# Analyses of rare and common alleles in parent-proband trios implicate rare missense variants in *SLC6A1* in schizophrenia and confirm the involvement of loss of function intolerant and neurodevelopmental disorder genes

**DOI:** 10.1101/607549

**Authors:** Elliott Rees, Jun Han, Joanne Morgan, Noa Carrera, Valentina Escott-Price, Andrew J. Pocklington, Madeleine Duffield, Lynsey Hall, Sophie E. Legge, Antonio F. Pardiñas, Alexander L. Richards, Julian Roth, Tatyana Lezheiko, Nikolay Kondratyev, Vera Golimbet, Mara Parellada, Javier González-Peñas, Celso Arango, GROUP Investigators, Micha Gawlik, George Kirov, James T. R. Walters, Peter Holmans, Michael C. O’Donovan, Michael J. Owen

## Abstract

Schizophrenia is a highly polygenic disorder with important contributions coming from both common and rare risk alleles, the latter including CNVs and rare coding variants (RCVs), sometimes occurring as *de novo* variants (DNVs). We performed DNV analysis in whole exome-sequencing data obtained from a new sample of 613 schizophrenia trios, and combined this with published data for a total of 3,444 trios. Loss-of-function (LoF) DNVs were significantly enriched among 3,488 LoF intolerant genes in our new trio data (rate ratio (RR) (95% CI) = 2.23 (1.31, 3.79); *p* = 2.2 × 10^−3^), supporting previous findings. In the full dataset, genes associated with neurodevelopmental disorders (NDD; n=160) were significantly enriched for LoF DNVs (RR (95% CI) = 3.32 (2.0, 5.21); *p* = 7.4 × 10^−6^). Within this set of NDD genes, *SLC6A1*, encoding a gamma-aminobutyric acid transporter, was associated with missense-damaging DNVs (*p* = 5.2 × 10^−5^). Using data from a subset of 1,122 trios for which we had genome-wide common variant data, schizophrenia polygenic risk was significantly over-transmitted to probands (*p* = 2.6 × 10^−60^), as was bipolar disorder common variant polygenic risk (*p* = 5.7 × 10^−17^). We defined carriers of candidate schizophrenia-related DNVs as those with LoF or deletion DNVs in LoF intolerant or NDD genes. These individuals had significantly less over-transmission of common risk alleles than non-carriers (*p* = 3.5 × 10^−4^), providing robust support for the hypothesis that this set of DNVs is enriched for those related to schizophrenia.

## Introduction

Genetic liability to schizophrenia involves a combination of rare and common risk alleles distributed across the genome^1^. Common schizophrenia risk alleles with odds ratios < 1.3 account for at least a third of genetic liability^2-4^, although only a small fraction of this is captured by the 145 genome-wide significant loci that were implicated in the largest published genome-wide association study (GWAS) of the disorder^5^. At the other end of the frequency spectrum, rare copy number variants (CNVs) and rare coding variants (RCVs), both sometimes occurring as *de novo* variants (DNVs), have been implicated in the disorder^6,7^. Although CNVs and RCVs are enriched in schizophrenia, not all rare variants observed in individuals with schizophrenia, including those occurring *de novo*, are expected to be aetiologically relevant, as there is a baseline burden of these variants in the general population.

In people with other neurodevelopmental disorders (NDDs) in which CNVs and RCVs play a role, particularly autism spectrum disorder (ASD)^8,9^ and developmental delay (DD)^10,11^, the enrichment for RCVs is greatest in genes classified as intolerant to loss-of-function (LoF) variants (i.e. variants that introduce premature stop codons or frameshifts in the encoded protein, or are predicted to disrupt mRNA splicing). This indicates that RCVs in these genes are more likely to be pathogenic for those disorders than RCVs occurring elsewhere in the genome. Moreover, greater enrichment is found for LoF DNVs than for missense DNVs that change an encoded amino acid, indicating the former class of mutation is particularly likely to be pathogenic. Similar observations have been made in schizophrenia, where an excess of LoF DNVs was found to be largely restricted to LoF intolerant genes^7^, although the degree of enrichment is lower than for ASD or DD.

In studies of ASD and DD, a significant excess of RCVs has been observed for 99 and 94 genes, respectively, with 32 of these genes overlapping between these disorders^8,10^. Only two genes, *SETD1A*^12^ and *RBM12*^13^, are currently associated with RCVs in schizophrenia. This is partly because of lower statistical power, as the number of trios that have been exome-sequenced in studies of schizophrenia (n=2,834) is smaller than equivalent studies of DD (n=7,580)^10^ and ASD (n = 6,430)^8^, but it also reflects the weaker enrichment in schizophrenia for this type of variant. As a set, genes disrupted by DNVs in NDDs are also enriched for DNVs in schizophrenia^14,15^, and therefore it follows that some of the genes implicated in ASD and DD by RCVs are also involved in the aetiology of schizophrenia. Aiming to contribute to the schizophrenia rare variant discovery effort, we have undertaken exome-sequencing in a new sample of 613 schizophrenia trios, and combined our data with published data from 2,834 trios, which includes 617 trios previously sequenced by our group^14^, to provide the largest analysis of coding DNVs in schizophrenia to date. Given the anticipated modest power even of this sample, as we have successfully done before for CNV analysis^16^, we exploited the well documented overlap in the genetic aetiologies of schizophrenia, ASD, and DD, to undertake a hypothesis focused analysis of NDD genes in schizophrenia which highlights *SLC6A1* as a novel risk gene.

The involvement of common variant polygenic risk in schizophrenia is already established^2,4,17^, but few existing studies have empirically examined the relationships between different classes of rare and common variants. In schizophrenia, the evidence indicates that rare CNVs and common risk variants co-act and do so in an additive manner^18,19^. Thus, affected carriers of schizophrenia-associated CNVs have been shown to have an elevated burden of common schizophrenia risk alleles as measured by the polygenic risk score (PRS), and this is inversely proportional to the estimated effect size of the implicated CNV^18^. Analogous observations have been made in ASD and DD, where common variant polygenic risk for those disorders has been shown to be over-transmitted from parents to probands, even in those that carry a disorder-associated DNV^20,21^. However, as yet, the relationship between rare exonic DNVs and common allele risk has not been studied in schizophrenia. Here, we examine this relationship using the polygenic transmission-disequilibrium test (pTDT)^21^. Specifically, we show that people with schizophrenia who are carriers of DNVs in gene sets proposed to be relevant to schizophrenia have a lower common risk allele burden than people with schizophrenia who are not carriers.

## Methods

### Sample overview

674 schizophrenia proband-parent trios, consisting of 2,000 individuals, were exome-sequenced on Illumina HiSeq 4000 platforms. The proband-parent trios were composed of 653 trios, 9 quads (two affected children) and one family with 3 affected children. None of these samples have been previously exome-sequenced. The proband-parent trios were ascertained from sites in Europe (Supplementary Table S1), each of which is described fully in the Sample Description section of the Supplementary Material. All probands had received a DSM-IV or ICD-10 diagnosis of schizophrenia.

### Exome sequence generation

Exome sequence was generated using the Nextera DNA Exome capture kit and HiSeq 3000/4000 PE Cluster Kit and HiSeq 3000/4000 SBS Kit. Raw sequencing reads were processed according to GATK best practice guidelines^22,23^. Reads were aligned to the human reference genome (GRCh37) using bwa version 0.7.15^24^. Variants were called using GATK haplotype caller (v3.4) and filtered using the GATK Variant Quality Score Recalibration (VQSR) tool. For all samples passing QC (criteria outlined below), we generated sequence data for a median of 83% of the exome target at ≥ 10X coverage.

### Sample quality control

Trios (n=27) were excluded for low sequencing coverage if less than 70% of the exome target achieved ≥10X coverage in the proband or either parent (Supplementary Figure S1). An additional 27 trios were excluded for excess heterozygosity (heterozygote:homozygote ratio > 1.9) or evidence of cross sample contamination (as measured by the FREEMIX sequence only estimate of contamination^25^) (Supplementary Figure S2). The last two metrics are highly correlated. Identity-by-descent (IBD) analysis (plink v1.9) to ensure expected proband-parent relationships resulted in exclusion of 3 trios. Four additional trios were excluded as outliers for the number of DNVs (Supplementary Figure S3). Following implementation of all the above sample QC steps, 613 proband-parent trios were retained for DNV analysis.

### Variant QC

In each of our newly sequenced samples, we excluded genotypes if they did not meet the following criteria: depth ≥ 10X; genotype quality score ≥ 30; allele balance ≤ 0.1 and ≥ 0.9 for homozygous genotypes for the reference and alternative allele, respectively; allele balance between 0.2 and 0.8 for heterozygous genotypes. For samples and variants that passed QC, we observed no difference in the number of heterozygous variants transmitted or non-transmitted from parents to probands (transmission disequilibrium test *p* = 0.53), indicating high data quality.

### *De novo* variant calling

Putative DNVs in the new trios were identified as sites that were heterozygous in the proband and homozygous for the reference allele in both parents. All trio members were required to pass genotype QC described above. We considered as putative DNVs 1) those where there were no reads for the mutant allele in either parent, and the mutant allele was not called in any other sample of the new trios (parent or proband) and 2) those where the mutant allele met all of the following; an allele count ≤3 in all newly sequenced samples, no mutant allele variant reads in either parent, and at least 5 reads of the mutant allele in the proband. Read alignments for all putative DNVs were manually inspected using IGV (http://software.broadinstitute.org/software/igv/) and variants were reassigned as high or low confidence if there was, respectively, no evidence or evidence for, read misalignment.

We used Sanger sequencing to perform a validation experiment, where DNA was available and primers could be designed, on all high confidence LoF DNVs, as well as additional putative DNVs. In total, primers were successfully designed for 205 putative DNVs. We observed high validation rates for high confidence DNVs (95.5%) and low rates (3.4%) for low confidence DNVs (Supplementary Table S2). Following these results, in our new trios we included in the downstream analyses all high confidence DNVs (N = 606 coding DNVs, Supplementary Table S3).

### Adding published de novo data

To increase the power of our analysis, we included previously published DNVs from 2,831 schizophrenia trios. When combined with our new trios, this resulted in a sample size of 3,444 schizophrenia trios. A summary of the published data can be found in Supplementary Table S4.

### De novo variant analysis

We tested whether DNVs were enriched in single genes or sets of genes using the statistical framework described in Samocha *et al* 2014^26^. Here, for a given set of genes we estimated the number of DNVs expected in our new sample using per-gene mutation rates^27^, adjusted for sequence coverage. When estimating the number of expected DNVs in previously published trios, we did not adjust per-gene mutation rates for coverage as coverage metrics were not available for all samples; the use of unadjusted per-gene mutation would over-estimate the expected number of DNVs in these trios, producing more conservative enrichment results. For single genes, we tested whether the overall burden of DNVs was significantly greater than that expected using a one-sided Poisson test (implemented in R). For our primary *de novo* gene set analysis, we controlled for background *de novo* rates by using a two-sample Poisson rate ratio test, which compared the DNV enrichment observed for genes in the set to that in genes outside the set.

DNVs were annotated using Ensemble Variant Effect Predictor (version 91)^28^. We define LoF variants as stop-gain, splice-acceptor, splice-donor and frameshift mutations. Although we observed a small number of start-loss and stop-loss DNVs, we did not include them in our LoF annotation as mutation rates are not available for these variants. We classify missense-damaging variants as missense variants with an MPC score ≥ 2, as this metric has proven effective at identifying variants associated with ASD^8,29^. Missense-damaging mutation rates for individual genes were calculated by summing tri-nucleotide mutation probabilities for all sites with an MPC score ≥ 2. Following previous work by us and others^14,15^, if an individual carried multiple *de novo* variants in the same gene, we conservatively considered these to be the result of a single mutation event, and retained for analysis only the variant predicted to be most deleterious.

### Polygenic risk scores

Where available (n=1,122 trios), we used SNP genotype data to generate polygenic risk scores. We confirmed that genotype and exome-sequence data belonged to the same individual through IBD analysis (plink v1.9). A summary of the data sets for which we had both exome-sequencing and SNP genotype data can be found in Supplementary Table S5. Using the largest available GWAS summary statistics that were independent from our trio test data, we derived PRS for schizophrenia, bipolar disorder (BD) and type 2 diabetes (T2D). We used BD PRS as previous studies have shown that common variant liability to schizophrenia and BD is substantially shared^30^. T2D PRS was used as a negative control. A summary of the training data used to generate PRS can be found in Supplementary Table S5.

For QC purposes, SNP genotype data were first harmonised to the Haplotype Reference Consortium panel using the Genotype Harmonizer package^31^ and then subjected to standard quality control, which included exclusion of samples with a call rate < 95%, SNPs with a MAF < 0.1, SNPs with > 1% missingness, or SNPs with a Hardy-Weinberg equilibrium exact test *p* value < 1 × 10^−6^. PRS were generated using PRSice 2 software^32^, where SNPs were clumped based on a window of 250 kb and a maximum r^2^ of 0.2. We generated PRS across a range of training data P-value thresholds (P < 0.5, 0.1, 0.05, 0.001).

### pTDT deviation

To test for a significant over-transmission of polygenic risk, we used the polygenic transmission disequilibrium test (pTDT) as described in Weiner *et al* (2017)^21^. Here, pTDT deviation scores were generated for each trio by subtracting the mean-parental PRS from the child PRS (Equation 1). pTDT deviation scores were standardised by dividing them by the cohort-specific mean-parental PRS standard deviation.

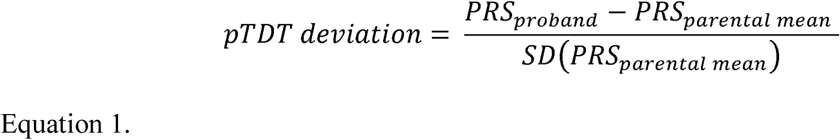

We tested whether the mean pTDT deviation was significantly greater than 0, representing an over-transmission of polygenic risk, by using a one-sided one-sample *t* test. A two-sample *t* test was used to compare mean pTDT deviation scores across groups of trios.

The primary pTDT results were produced using PRS generated with a *P*-threshold of 0.05, as this threshold explained the most case-control variance in the 2014 schizophrenia PGC analysis^4^. However, we also present in the Supplementary Material Table S6 pTDT results obtained for PRS generated across different *P*-value thresholds.

## Results

### De novo mutation rates

After variant QC, we observed 606 coding *de novo* variants (DNVs) in 613 probands (Supplementary Table S3), corresponding to a rate of 0.99 (s.e = 0.041) events per proband, which is not significantly different to the rate observed in a sample of 617 schizophrenia trios previously published by our group^14^ (previous *de novo* rate = 1.032; rate ratio (RR) (95% confidence interval (CI)) = 0.96 (0.86, 1.07); *p* = 0.46). Of these coding DNVs, 154 were synonymous, 372 were missense, 15 were inframe indels, 2 start-loss, 1 stop-lost, and 62 were LoF (19 stop-gain, 13 splice and 30 frameshift indels). The number of coding DNVs observed per-trio followed the expected Poisson distribution (Supplementary Figure S4).

### De novo variant enrichment tests

In the new data set, we observed a significant excess of LoF DNVs among LoF intolerant genes (Fig 1, RR (95% CI) = 2.23 (1.31, 3.79); *p* = 2.2 × 10^−3^; Supplementary Table S7). Consistent with previous reports, we found no evidence for DNV enrichment in the following negative control gene set tests: LoF DNVs in LoF tolerant genes (Fig 1), synonymous DNVs in LoF intolerant genes; synonymous DNVs in LoF tolerant genes (Supplementary Table S8). After combining the new trio data with previously published data from 2,831 trios, LoF DNVs were enriched in LoF intolerant genes with a RR (95% CI) of 1.58 (1.28, 1.96) (Fig 1, *p* = 2.5 × 10^−5^; Supplementary Table S7).

**Figure 1.**
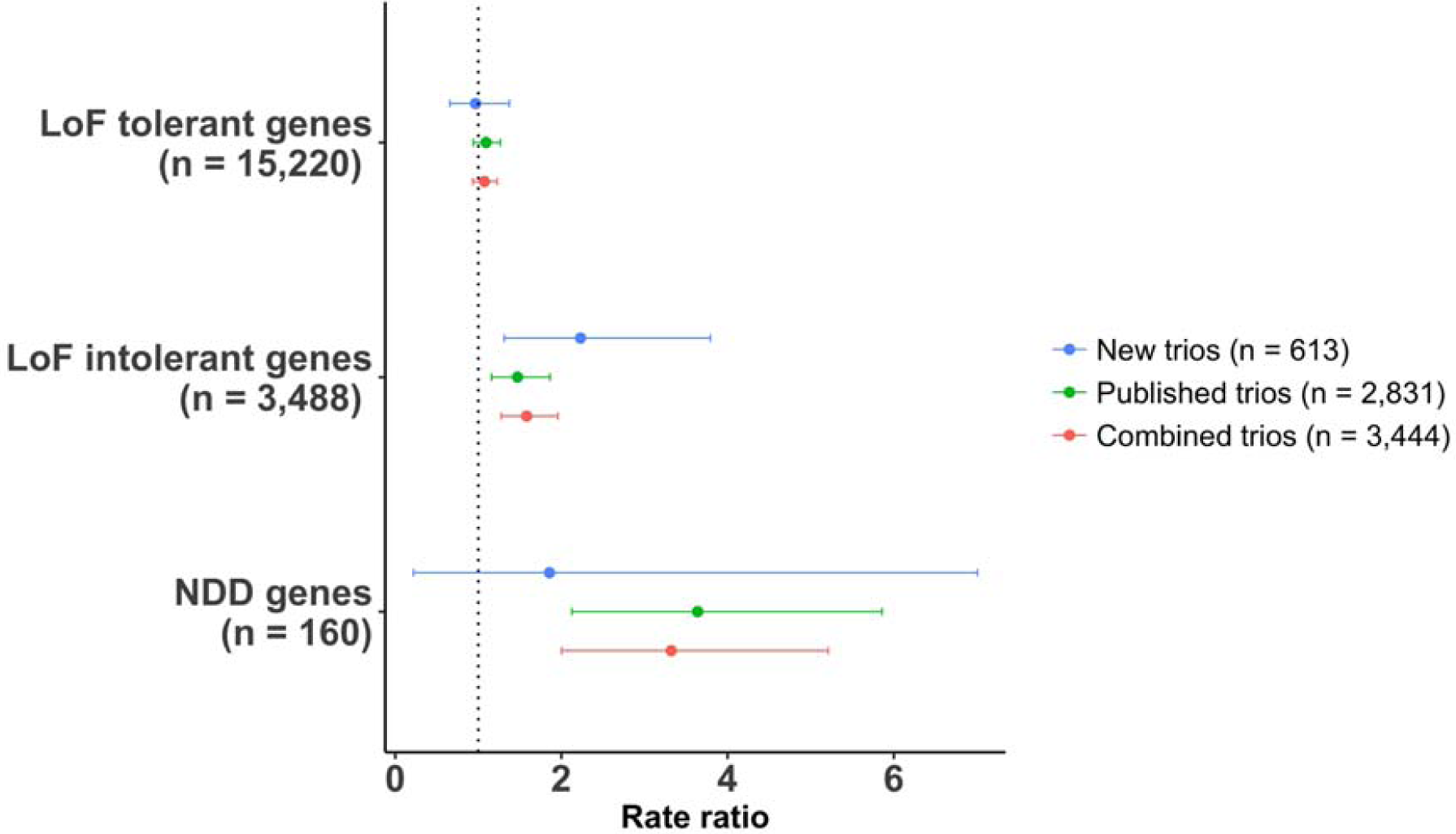
Gene set enrichment for loss-of-function *de novo* variants. Loss-of-function (LoF) DNVs were tested in LoF intolerant genes and neurodevelopmental disorder (NDD) genes. For LoF intolerant and NDD gene sets, rate ratios and 95% confidence intervals are relative to the baseline DNV rate, which is defined as the LoF DNV enrichment observed for all genes outside of the given set. LoF DNV enrichment for LoF tolerant genes are shown as a negative control. A breakdown of the LoF intolerant and NDD gene set results is provided in Supplementary Tables S7 and S8.

In the combined trio data, no individual gene was significantly enriched for LoF DNVs after correction for all genes tested (n=18,658). The most significant novel gene was *CUL1*, which had two LoF DNVs in the new trios and one additional LoF DNV in the published trios (Table 1).

**Table 1.** Genes disrupted by 2 or more LoF *de novo* variants. The most significant gene, *SETD1A*, has been previously identified as a schizophrenia risk gene^12^.

| Gene | New trios<br>(n=613) |  | Published trios<br>(n=2,831) |  | All trios<br>(n=3,444) |  |
| --- | --- | --- | --- | --- | --- | --- |
|  | LoF DNVs | P | LoF DNVs | P | LoF DNVs | P |
| *SETD1A* | 0 | 1 | 3 | 1.90E-06 | 3 | 3.00E-06 |
| *CUL1* | 2 | 3.60E-05 | 1 | 0.04 | 3 | 2.00E-05 |
| *TAF13* | 0 | 1 | 2 | 2.40E-05 | 2 | 3.40E-05 |
| *GALNT9* | 0 | 1 | 2 | 2.90E-05 | 2 | 4.20E-05 |
| *HENMT1* | 0 | 1 | 2 | 5.50E-05 | 2 | 7.90E-05 |
| *PAF1* | 1 | 0.0028 | 1 | 0.013 | 2 | 0.00013 |
| *SV2B* | 0 | 1 | 2 | 0.00016 | 2 | 0.00023 |
| *NRXN3* | 0 | 1 | 2 | 0.00022 | 2 | 0.0003 |
| *HIVEP3* | 0 | 1 | 2 | 0.00026 | 2 | 0.00035 |
| *RB1CC1* | 0 | 1 | 2 | 0.00046 | 2 | 0.00064 |
| *SMARCC2* | 0 | 1 | 2 | 0.0005 | 2 | 0.00068 |
| *MKI67* | 0 | 1 | 2 | 0.00085 | 2 | 0.0012 |
| *CHD8* | 0 | 1 | 2 | 0.0009 | 2 | 0.0013 |
| *TENM1* | 1 | 0.0077 | 1 | 0.046 | 2 | 0.0014 |
| *TRIO* | 0 | 1 | 2 | 0.0012 | 2 | 0.0016 |
| *SCN2A* | 1 | 0.012 | 1 | 0.057 | 2 | 0.0024 |
| *DNAH9* | 0 | 1 | 2 | 0.0018 | 2 | 0.0026 |
| *KMT2C* | 0 | 1 | 2 | 0.0086 | 2 | 0.012 |
| *KIAA1109* | 0 | 1 | 2 | 0.01 | 2 | 0.015 |
| *TTN* | 1 | 0.16 | 2 | 0.22 | 3 | 0.093 |

We have previously shown that rare CNVs that increase risk of schizophrenia are effectively confined to those that also influence other NDDs^16^. Defining NDD genes as those (N=160) that are significantly enriched for RCVs in recent large studies of ASD^8^ and DD^10^, NDD genes were significantly enriched for LoF DNVs in the combined trio data (Fig 1; RR (95% CI) = 3.32 (2.0, 5.21); *p* = 7.4 × 10^−6^; Supplementary Table S7), and this enrichment was significantly greater than for LoF intolerant genes (RR (95% CI) = 2.37 (1.41, 3.8); *p* = 8.6 × 10^−4^).

Exploiting the strong enrichment among NDD genes for DNVs in schizophrenia, we undertook focused analysis of genes in this set, with the aim of identifying high probability schizophrenia risk genes. As highlighted in the study of ASD^8^, association to some NDD genes is driven by LoF variants alone, a combination of LoF variants and missense variants, and in some cases, primarily by missense variants. Therefore, we considered all those classes of mutation in our analysis. All LoF/missense-damaging DNVs observed in NDD genes and, where available, phenotypes observed in these carriers are presented in Supplementary Table S9.

*SLC6A1* was significantly associated with missense-damaging DNVs in our new trio data after correcting for three classes of mutation (LoF, missense-damaging and LoF plus missense-damaging) and 160 NDD genes (2 damaging-missense DNVs; *p* = 7.46 × 10^−5^; *p*_*corrected*_ = 0.036). This finding was supported in our analysis of all trio data, where we observed one additional missense-damaging DNV (Table 2; 3 missense-damaging DNVs; *p* = 5.2 × 10^−5^, *p*_*corrected*_ = 0.025). It is striking that in the study of ASD^8^, association to *SLC6A1* was also driven by missense variants (n=8) rather than LoF variants (n=1). Following the rationale outlined by the Deciphering Developmental Disorders Study^33^, we undertook a combined analysis of schizophrenia and ASD DNVs; the evidence for enrichment of missense-damaging DNVs (MPC ≥ 2) in *SLC6A1* was more than 3 orders of magnitude stronger than for ASD alone, supporting the hypothesis that missense variants in this gene contribute to both disorders (combined *p* = 1.6 × 10^−14^; ASD alone *p* = 8.0 × 10^−11^).

**Table 2.**
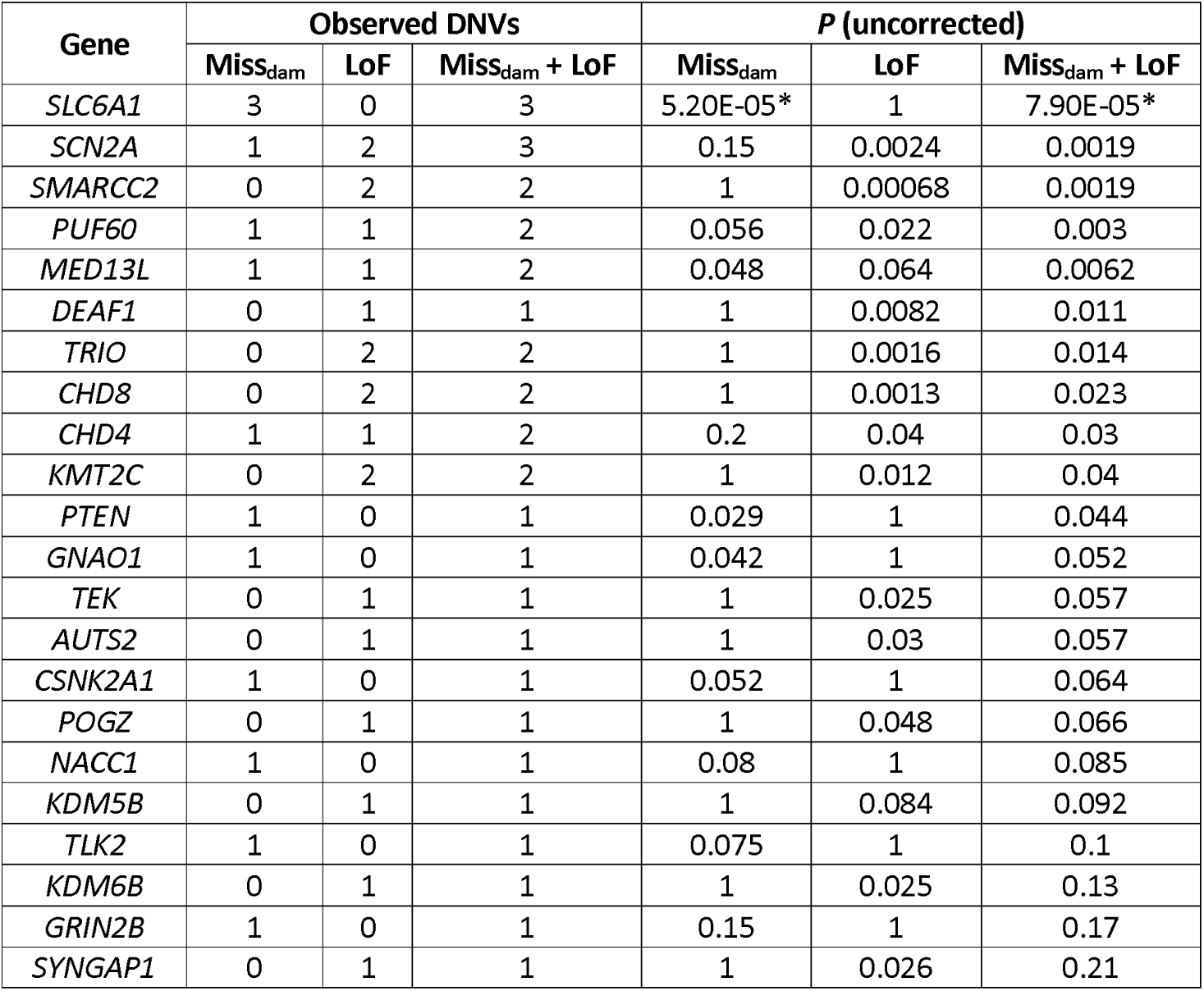
Neurodevelopmental disorder genes with at least 1 LoF or missense-damaging *de novo* variant observed in schizophrenia. Enrichment *P* values are derived from the analysis of all schizophrenia trios (n = 3,444). Miss_dam_ = missense-damaging (MPC score ≥ 2). * indicate *p* values which survive correction for 160 NDD genes and three mutation classes (LoF, missense-damaging and LoF plus missense-damaging).

| Gene | Observed DNVs |  |  | *P* (uncorrected) |  |  |
| --- | --- | --- | --- | --- | --- | --- |
|  | Miss<sub>dam</sub> | LoF | Miss<sub>dam</sub> + LoF | Miss<sub>dam</sub> | LoF | Miss<sub>dam</sub> + LoF |
| *SLC6A1* | 3 | 0 | 3 | 5.20E-05* | 1 | 7.90E-05* |
| *SCN2A* | 1 | 2 | 3 | 0.15 | 0.0024 | 0.0019 |
| *SMARCC2* | 0 | 2 | 2 | 1 | 0.00068 | 0.0019 |
| *PUF60* | 1 | 1 | 2 | 0.056 | 0.022 | 0.003 |
| *MED13L* | 1 | 1 | 2 | 0.048 | 0.064 | 0.0062 |
| *DEAF1* | 0 | 1 | 1 | 1 | 0.0082 | 0.011 |
| *TRIO* | 0 | 2 | 2 | 1 | 0.0016 | 0.014 |
| *CHD8* | 0 | 2 | 2 | 1 | 0.0013 | 0.023 |
| *CHD4* | 1 | 1 | 2 | 0.2 | 0.04 | 0.03 |
| *KMT2C* | 0 | 2 | 2 | 1 | 0.012 | 0.04 |
| *PTEN* | 1 | 0 | 1 | 0.029 | 1 | 0.044 |
| *GNAO1* | 1 | 0 | 1 | 0.042 | 1 | 0.052 |
| *TEK* | 0 | 1 | 1 | 1 | 0.025 | 0.057 |
| *AUTS2* | 0 | 1 | 1 | 1 | 0.03 | 0.057 |
| *CSNK2A1* | 1 | 0 | 1 | 0.052 | 1 | 0.064 |
| *POGZ* | 0 | 1 | 1 | 1 | 0.048 | 0.066 |
| *NACC1* | 1 | 0 | 1 | 0.08 | 1 | 0.085 |
| *KDM5B* | 0 | 1 | 1 | 1 | 0.084 | 0.092 |
| *TLK2* | 1 | 0 | 1 | 0.075 | 1 | 0.1 |
| *KDM6B* | 0 | 1 | 1 | 1 | 0.025 | 0.13 |
| *GRIN2B* | 1 | 0 | 1 | 0.15 | 1 | 0.17 |
| *SYNGAP1* | 0 | 1 | 1 | 1 | 0.026 | 0.21 |

### Polygenic transmission disequilibrium tests

Schizophrenia and BD PRS were significantly over-transmitted from parents to probands (Fig 2, Supplementary Material Table S6). Under a liability threshold model, probands carrying DNVs of large effect size should require less transmission of polygenic risk than probands without such a variant. To test this, the mean pTDT was compared between carriers of candidate schizophrenia related DNVs and the remainder of the sample. Following the results of the gene set analysis, we define candidate schizophrenia related DNVs as LoF DNVs in a LoF intolerant gene or a NDD gene. Given CNV deletions disrupting LoF intolerant genes are associated with schizophrenia^7^, we also included *de novo* CNV deletions disrupting one of these genes as candidate schizophrenia related DNVs (CNVs contributing to this analysis are presented in Supplementary Table S10. CNV calling procedure is outlined in the CNV section of the Supplementary Material).

**Figure 2.**
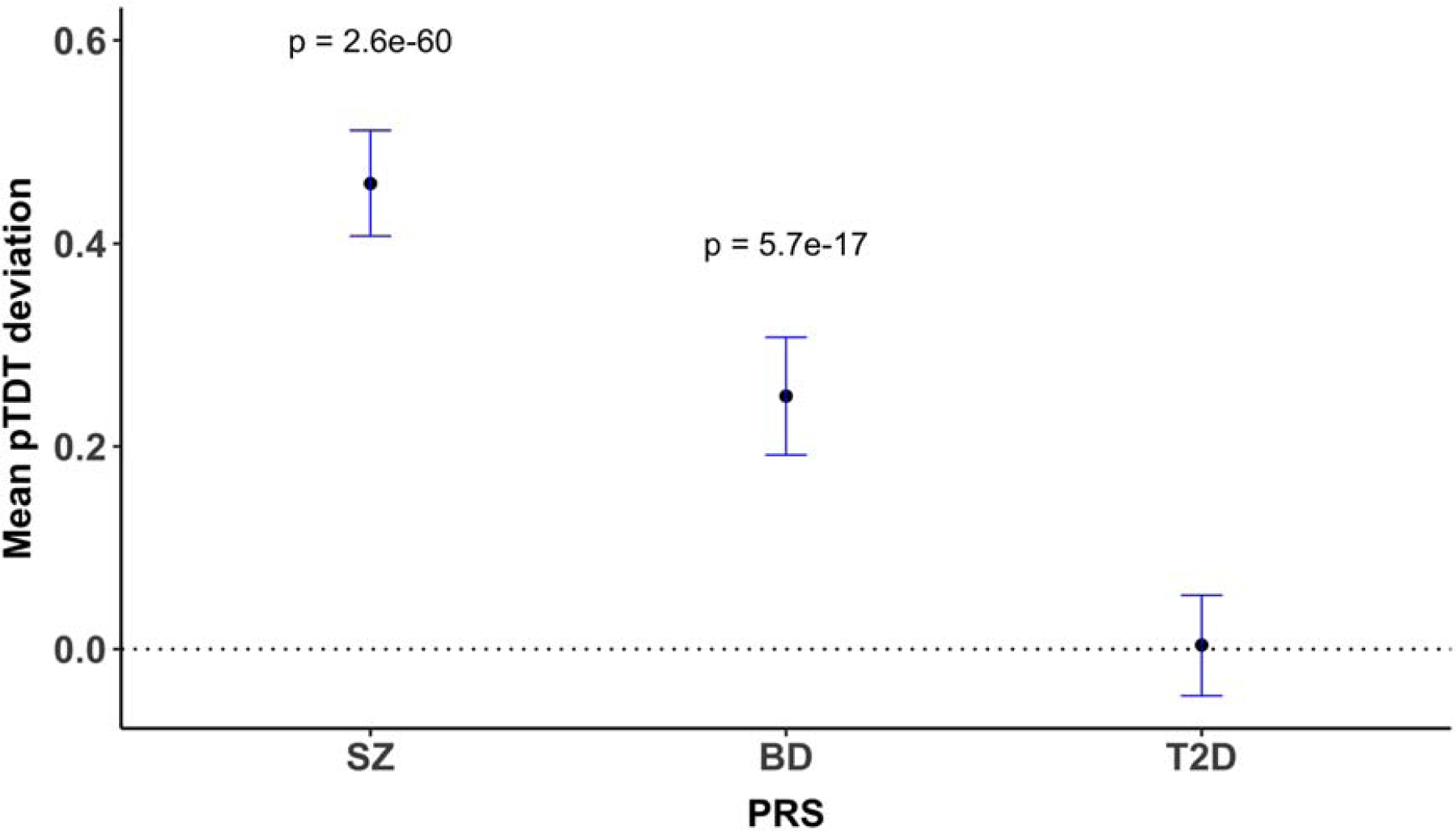
Mean pTDT deviation and 95% confidence intervals for schizophrenia, bipolar disorder (BD), and type 2 diabetes (T2D) PRS. Polygenic risk for schizophrenia and bipolar disorder is significantly over-transmitted to schizophrenia probands.

Probands carrying candidate schizophrenia related DNVs had a significantly lower mean pTDT than those who did not carry one of these DNVs (carrier mean pTDT (95% CI) = 0.07 (−0.15, 0.29); non-carrier mean pTDT (95% CI) = 0.48 (0.43, 0.54); *p* = 3.5 × 10^−4^; Fig 3). Based on mean pTDT point estimates, the over-transmission of common risk alleles from parents is about 7-fold greater to non-carriers than carriers of candidate schizophrenia related DNVs, although this estimate is imprecise given the width of the confidence intervals (Fig 3). Similar patterns were observed when LoF and deletion DNVs were tested separately (Fig 3). In a negative control test, the mean pTDT did not significantly differ between probands carrying a synonymous DNV in either a LoF intolerant or NDD gene and non-carriers (Fig 3).

**Figure 3.**
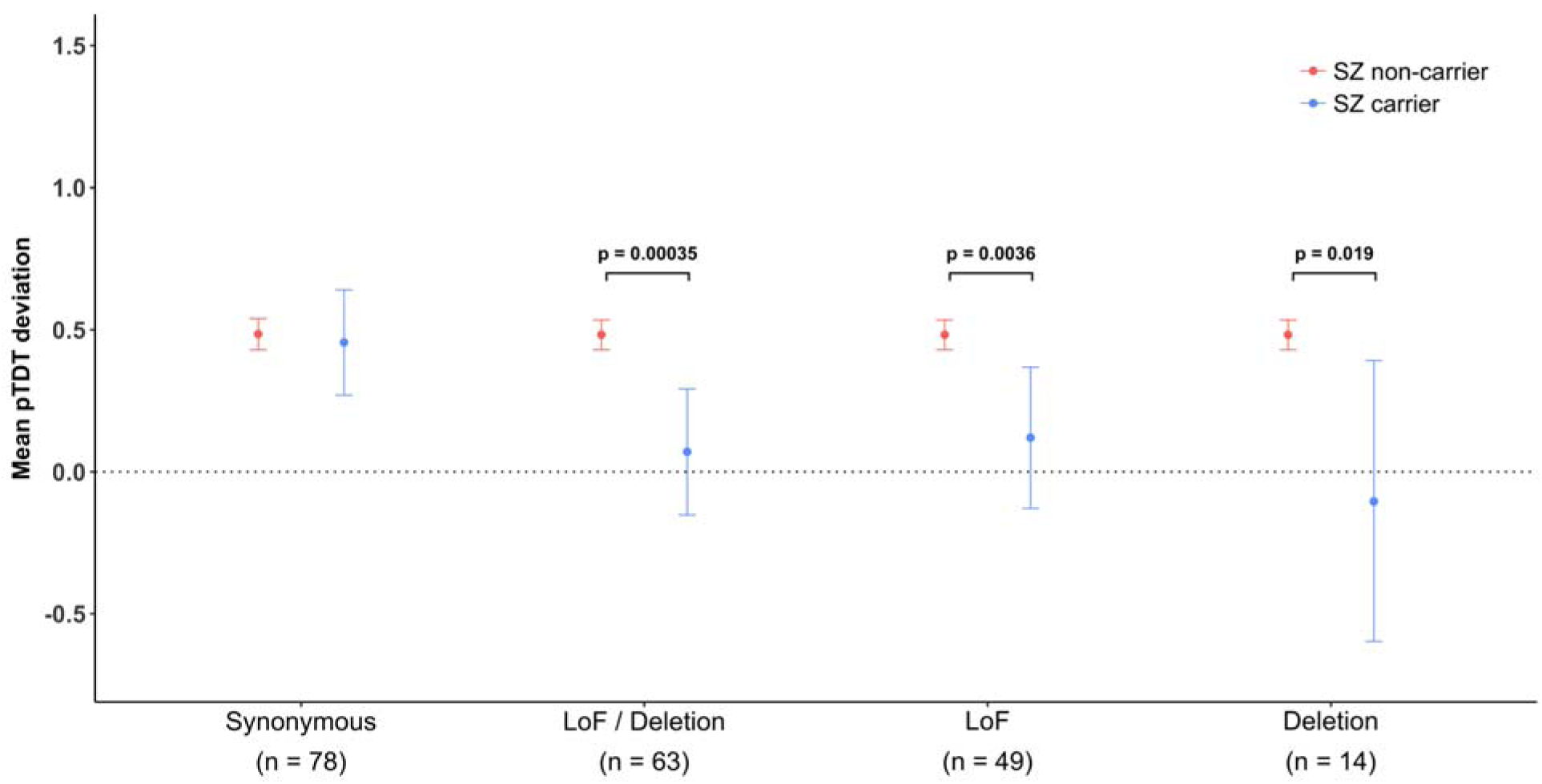
Mean pTDT deviation and 95% confidence intervals for schizophrenia PRS. Results are shown for probands carrying various classes of *de novo* variant (DNV) in a LoF intolerant gene or a neurodevelopmental disorder gene; our primary analysis defined schizophrenia carriers as probands with a LoF or deletion DNV in a LoF intolerant gene or a neurodevelopmental disorder gene (LoF/deletion label). Results are also shown separately for carriers of LoF and deletion DNVs. LoF = loss-of-function.

The finding that the mean pTDT deviation for schizophrenia PRS was significantly different between probands carrying candidate schizophrenia related DNVs and non-carrying probands was consistent across schizophrenia PRS training *p*-value thresholds (Supplementary Table S11). Although the pTDT method is expected to be robust to population stratification, the efficacy of PRS as a measure of relative liability varies with the extent to which the ancestry of the sample from which risk alleles are derived (the source GWAS) matches the ancestry of those being tested (in our case the trios). Given the source GWAS is primarily of European ancestry, we tested, and confirmed, that our findings held when we restrict our analysis to trios with European ancestry (as defined by PCA; Supplementary Figure S5) despite the smaller sample size (Supplementary Table S11).

The mean pTDT in carriers of candidate schizophrenia related DNVs was not significantly greater than the null (Fig 3). Based on the pTDT standard deviation observed for schizophrenia PRS in all trios (0.89), we only had 80% power to detect a significant (alpha = 0.05) mean pTDT of 0.4 in the 63 carriers of candidate schizophrenia related DNVs. Thus, while we can be confident that the over-transmission to candidate DNV carriers is less than to carriers, power limitations mean we cannot conclude that candidate DNV carriers have no contribution from common alleles.

## Discussion

Proband-parent trio studies have identified large numbers of genes associated with DNVs in ASD and DD^8,10^. Although similar studies in schizophrenia have revealed general pathophysiological insights into the disorder, such as a role in the disorder for proteins involved in postsynaptic signaling complexes^14,34^, schizophrenia gene discovery through DNV analysis has been hindered by small samples. To add to efforts to overcome this limitation, we performed exome-sequencing of a new sample of 613 schizophrenia trios. We confirmed previous work showing schizophrenia LoF DNVs are significantly enriched among a set of 3,488 genes intolerant to this class of mutation, and identified a stronger enrichment of DNVs in a smaller set of 160 genes that are associated with RCVs in NDDs.

In our analysis of all schizophrenia trios, no novel gene was unequivocally associated with DNVs after correction for all genes tested. However, taking an approach based on the wealth of data showing that rare CNVs that increase risk of schizophrenia are effectively confined to those that also influence other NDDs^16^, and exploiting the observation here for strong enrichment for DNVs in known NDD genes, we find evidence for association between *SLC6A1*, which encodes a sodium-dependent γ-aminobutyric acid (GABA) transporter (also known as GAT1), and missense-damaging DNVs. *SLC6A1* is involved in reuptake of the inhibitory neurotransmitter GABA from the synaptic cleft; our finding therefore adds to the evidence for perturbation of GABAergic neuronal signaling in genetic risk for schizophrenia^35^. Congruent with our findings, the largest study of RCVs in ASD found *SLC6A1* to be the most significant (of only four) genes where association signal was driven by missense-damaging variants (8 missense and 1 LoF DNVs)^8^. In myoclonic atonic epilepsy and DD, LoF variants account for 54% and 30% of the observed nonsynonymous DNVs, respectively^8^. Given the strong convergent evidence for this gene, and specifically for a role for missense mutations, from other NDDs, *SLC6A1* is highly likely also to be involved in schizophrenia. This conclusion is further supported by the result of the DNV missense meta-analysis of ASD and schizophrenia, in which the combined evidence for association is more than 3 orders of magnitude stronger than the (already strong) evidence for association to ASD alone, and surpasses genome-wide significance by 8 orders of magnitude.

The role of polygenic risk in schizophrenia has been widely studied using large case-control samples. However, to our knowledge, this is the first study to investigate polygenic risk in schizophrenia using the pTDT method. The pTDT method has several advantages over case-control PRS studies as it is not confounded by ancestry or ascertainment bias or the possibility of effects arising from super-healthy controls in discovery GWAS and subsequent PRS test samples^21^. Our results provide strong refutation that such effects might explain the PRS effects that have been widely publicised in the literature, including that of overlap in risk between schizophrenia and BD. More importantly in the present context, our finding that carriers of LoF DNVs in the large gene set defined by LoF intolerance, or in a known NDD gene, have significantly lower distortion of transmission of polygenic liability from the mean parental PRS than do non-carriers provides orthogonal evidence that a substantial proportion of this class of variants contribute to schizophrenia pathogenesis. This is an important observation given the possibility that previously documented gene set enrichments in cases of these variants could have been driven by errors in the calibration of the expected mutation rate, or technical issues arising from comparing cases and controls (or case and control trios) often derived opportunistically from different studies.

In terms of magnitude of difference in the transmission distortion between probands carrying candidate schizophrenia related DNVs and those that do not, our limited sample size does not allow this to be accurately estimated. The point estimate is that the distortion in non-carriers is about 7-fold of that of carriers (and almost 10-fold when restricted to those of European ancestry). This suggests that on average, the candidate DNVs contribute substantially to liability in those who carry them, indeed carriers of candidate schizophrenia related DNVs did not significantly over-inherit schizophrenia PRS when compared with chance. However, it is important to stress that the latter finding most likely reflects limited power rather than no role for common variation in the carriers; consistent with this interpretation, the point estimate for the pTDT in candidate DNV carriers is greater than 0. Nevertheless, it will be important for future work to determine whether differences in co-action between common and rare risk alleles exist between schizophrenia and NDDs.

In conclusion, we provide further evidence that certain classes of DNV are associated with increased risk for schizophrenia. We highlight strong evidence that mutations in *SLC6A1*, a known ASD, DD and epilepsy gene, confer high risk of schizophrenia. Through combining exome-sequencing and GWAS data, we show that carriers of candidate schizophrenia related DNVs inherit significantly fewer common risk alleles than non-carrying cases, providing strong orthogonal evidence that these DNVs contribute to schizophrenia liability.

## Acknowledgments and Disclosures

The work at Cardiff University was supported by Medical Research Council Centre Grant No. MR/L010305/1 (to MJO) and Program Grant No. G0800509 (to MJO, MCO, JTRW, VE-P, PH, AJP), European Community Seventh Framework Programme Grant No. HEALTH-F2-2010-241909 (Project EU-GEI), and European Union Seventh Framework Programme for research, technological development, and demonstration Grant No. 279227 (CRESTAR Consortium). We acknowledge Lesley Bates and Lucinda Hopkins, at Cardiff University, for laboratory sample management. We acknowledge Mark Einon, at Cardiff University, for support with the use and setup of computational infrastructures.

## Competing interests

The authors declare no competing interests.

