## Supplemental material for "Analyses of rare and common alleles in parent-proband trios implicate rare missense variants in *SLC6A1* in schizophrenia and confirm the involvement of loss of function intolerant and neurodevelopmental disorder genes"

**Supplementary Material**

Ethics statement

All research conducted as part of this study was approved by ethical bodies and consistent with regulatory and ethical guidelines.

Sample Description

Bulgarian trios

We sequenced 77 proband-parent trios recruited from Bulgaria whose ascertainment and diagnosis are as described previously^1^. These trios were independent from a previous exome-sequencing study of Bulgarian samples from our group^2^. Briefly, all cases had been hospitalised and met DSM-IV criteria ^3^ for schizophrenia or schizoaffective disorder based upon SCAN (Schedules for Clinical Assessment in Neuropsychiatry)^4^ interview by psychiatrists, and review of case notes. Cases were recruited from general adult psychiatric services and were typical of those attending those services. All participants provided informed consent.

German trios

The German sample included 337 parent-proband trios. Patients were identified through hospital records or during inpatient stays or outpatient clinics. All research subjects and, where applicable, their legal guardians provided a written informed consent to participate in the study. The ethical committee of Wuerzburg reviewed and approved the study. Patients were diagnosed according to ICD-10 criteria, whereby a consensus diagnosis was made by at least two independent, trained raters based on all available clinical information standardized by the AMDP-System (Manual for Assessment and Documentation of Psychopathology in Psychiatry). DNA samples of the participants were extracted from peripheral blood.

Russian trios

The sample included 83 trios. Probands were inpatients at the psychiatric units of the Mental Health Research Centre, Moscow, Russia. All patients were diagnosed with schizophrenia or schizoaffective disorder. The diagnosis was made by two psychiatrists according to diagnostic criteria of ICD-10 and was based on medical records and a semi-structured interview (MINI, SADS). Interviews were conducted by trained researchers. All participants provided a written informed consent to molecular-genetic research. DNA was extracted from peripheral blood.

Spanish trios

The Spanish sample included 37 schizophrenia trios. Patients were diagnosed at the Hospital Gregorio Marañón, and were diagnosed with Schizophrenia or Schizophreniform disorder. Diagnoses were determined by clinical psychiatrists or psychologists, according to DSM-IV criteria with the Structured Clinical Interview for DSM I and II (SCID-I and II) for adults, and the Kiddie-Schedule for Affective Disorders & Schizophrenia, Present & Lifetime Version (K-SADS-PL) for participants aged under 18 years. The diagnostic interviews were administered both at baseline and at 2-years follow-up. DNA was extracted from peripheral blood.

UK trios

The schizophrenia families from the UK were recruited as part of sib-pair and case–control collections. This cohort has been described in detail elsewhere^5^. All probands had received a DSM-IV diagnosis of schizophrenia or schizoaffective disorder, where a consensus diagnosis was made by two independent, trained raters based on all available clinical information including a semi-structured interview [PSE-9 or Assessment of Symptoms and History or Schedules for Clinical Assessment for Neuropsychiatry (SCAN)^4^], examination of case notes and information from relatives and mental health professionals. All interviews were conducted by psychiatrists and psychologists after written consent was obtained following local ethical approval guidelines.

GROUP trios

The 91 GROUP families were recruited from several sites across the Netherlands. Cases were between 16 and 50 years of age, and had received a diagnosis of schizophrenia according to DSM-IV criteria. To assess DSM-IV diagnosis, the Comprehensive Assessment of Symptoms and History (CASH)^6^ or SCAN interviews^4^ were used. The study protocol was approved centrally by the Ethical Review Board of the University Medical Centre Utrecht and subsequently by local review boards of each participating institute. A detailed description of the GROUP cohort can be found here in Korver et al 2012^7^.

Supplementary figures

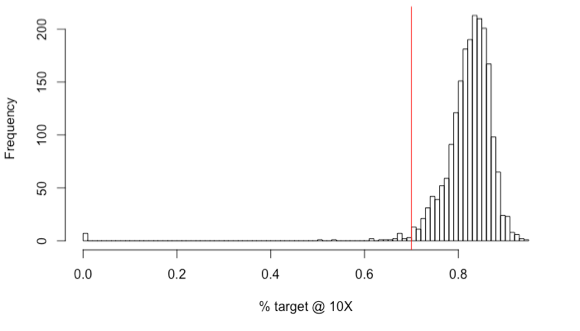

**Figure S1**. Proportion of exome target sequenced to ≥10X coverage in each sample. Red line indicates our cut off (70% of exome target) for excluding samples due to low coverage.

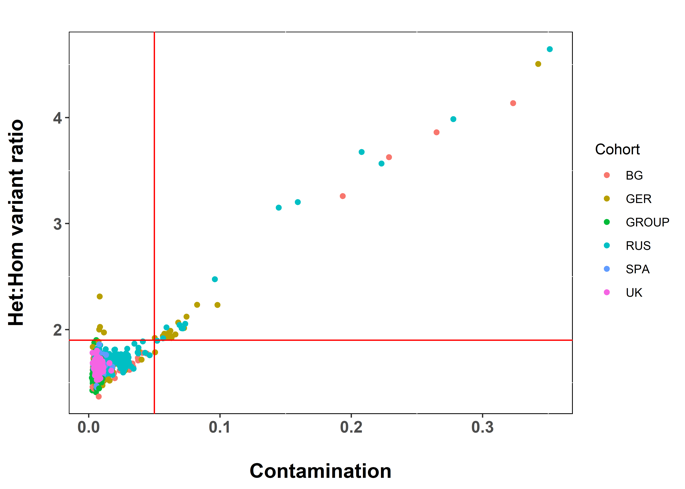

**Figure S2.** Exclusion thresholds for contamination and/or heterozygosity. Contamination was estimated using the FREEMIX sequence only estimate of contamination method^8^.

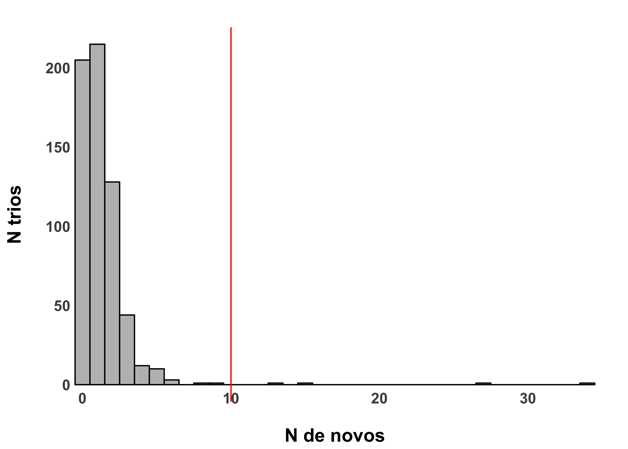

**Figure S3.** Number of *de novo* variants per trio. Red vertical line indicates the threshold used to exclude probands as outliers for number of *de novo* variants.

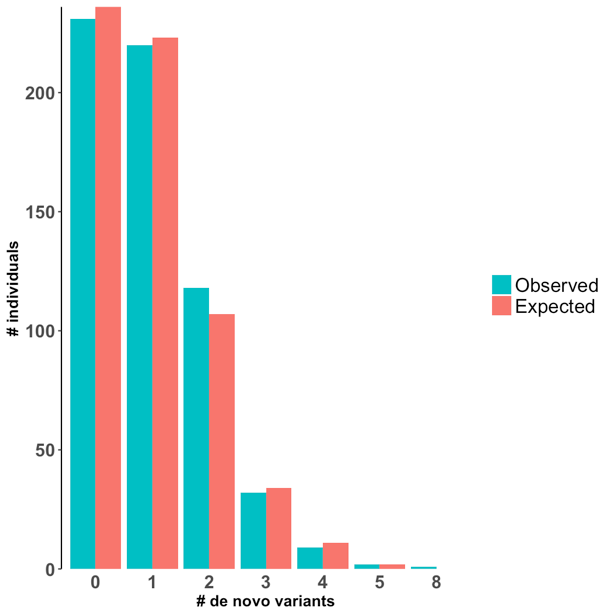

**Figure S4**. Distribution of number of coding variants per trio.

CNV analysis

For the German, Spanish, GROUP and Russian cohorts from the new trio sample, we used Log R ratio and B-allele frequency information from the SNP genotyping data to call CNVs using the PennCNV algorithm, following standard protocol and adjusting for GC content^9^. We performed our standard pipeline for CNV quality control^10^ to exclude samples that were outliers for LRR standard deviation, B-allele frequency drift, wave factor and total number of CNVs called per person. Individual CNV calls were excluded if they were < 10kb, covered by < 10 probes, > 50% of their length overlapped low copy repeats or the CNV was observed in more than 1% of the sample. The raw intensity traces for all putative *de novo* CNVs, defined as CNVs observed in the proband and no overlapping CNV in either parent, were manually inspected for data quality.

For the Bulgarian trios, *de novo* CNVs have been previously published by our group^1^.

Identifying sample ancestry through principle component analysis

We performed a principle component (PC) analysis using SNP genotype data from the 1000 genomes project, and then projected our proband-parent trios onto these PCs using EIGENSOFT smartPCA^11^. We did this separately for samples genotyped on Illumina psychchips (Supplementary Figure S5 A) and Affymetrix 6.0 arrays (Supplementary Figure S5 B), noting that all members of a trio were genotyped on the same array. For the purpose of excluding trios with non-European ancestry in the polygenic-transmission disequilibrium analysis, we defined European samples as those falling within the box surrounded by red lines shown in Supplementary Figure S5. For samples contributing to the pTDT analysis, this resulted in 128/1122 (11.4%) trios being excluded for having non-European ancestry.

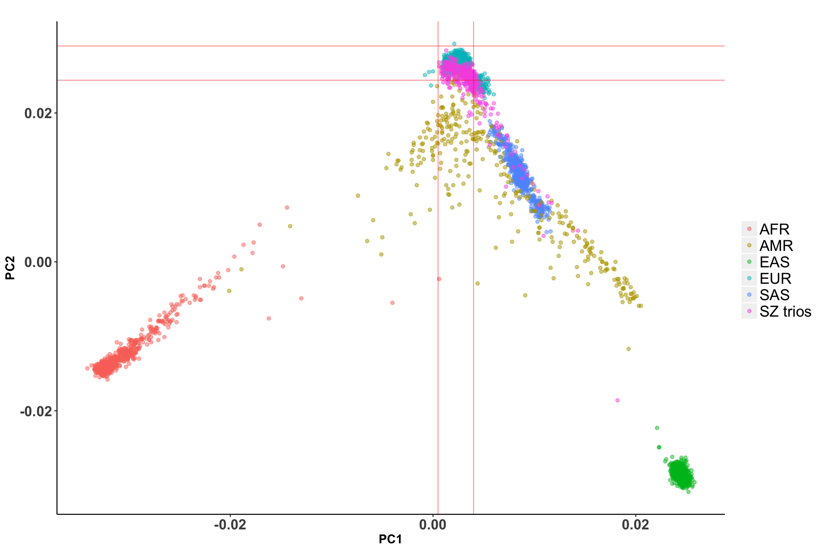

**A**

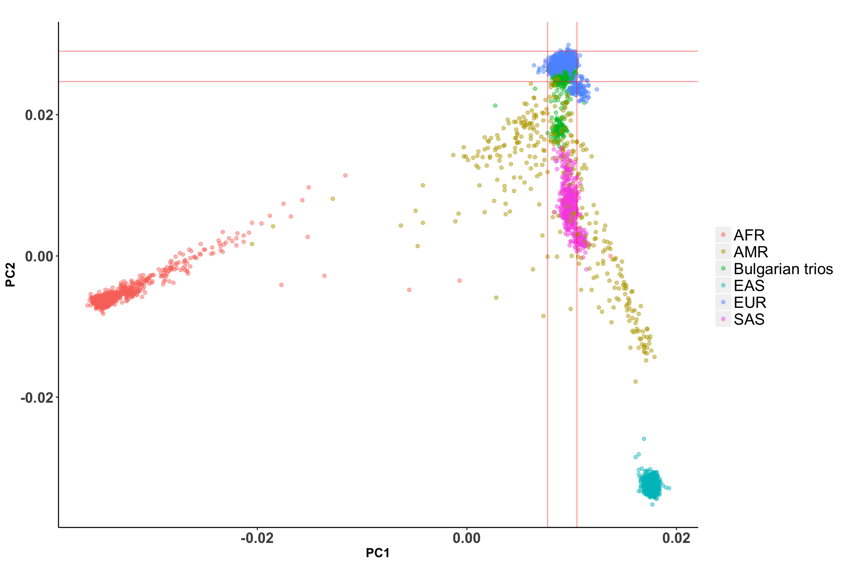

**B**

**Figure S5**. Sample ancestry. **A)** Samples contributing to proband-parent trios genotyped on Illumina psychchips. **B)** Samples contributing to proband-parent trios genotyped on Affymetrix 6.0 arrays, all of which were members of the Bulgarian trios.
